# The YTHDF1-3 proteins are bidirectionally influenced by the codon content of their mRNA targets

**DOI:** 10.1101/2023.11.20.565808

**Authors:** Clara Moch, Limei Zou, Nicolas Pythoud, Emilie Fillon, Gabrielle Bourgeois, Marc Graille, Christine Carapito, Clément Chapat

## Abstract

*N*^6^-methyladenosine (m^6^A) is the most abundant modification in eukaryotic mRNAs and plays critical roles in a broad variety of biological processes. Recognition of m^6^A by the YTHDF1-3 proteins results in the alteration of the translation efficiency and stability of methylated mRNAs, although their mode of action is still matter of debates. To decode the molecular basis of YTHDF1-3 action in human cells, we performed an unbiased proteomic screen of their full spectrum of interacting proteins using BioID (proximity-dependent biotin identification). Our systematic BioID mapping revealed that each YTHDF protein is a dynamic hub that associates with both mRNA silencing machineries and the translation apparatus. Based on this, we identified a direct interaction between YTHDF2 and the ribosomal protein RACK1, and found that the silencing activity of YTHDF2 is bidirectionally modulated by the codon content of its targeted mRNAs. Using a tethering reporter system that recapitulates this phenomenon, we confirmed that the three YTHDF proteins selectively repress mRNAs enriched in optimal codons, while they activate those enriched in non-optimal codons. Altogether, these results have important implications for understanding the underlying multiplicity of YTHDF1-3 and could reconcile seemingly contradictory data.

## INTRODUCTION

Post-transcriptional silencing mechanisms modulate global and transcript-specific mRNA stability and translation, contributing to the rapid and flexible control of gene expression. mRNA silencing relies on a variety of non-coding RNAs, such as microRNAs (miRNAs), mRNA modifications and RNA-binding proteins (RBPs) which target the 3’-untranslated regions (3’UTRs) of specific mRNAs to recruit silencing effector machineries. Among these modifications, *N*^6^-methyladenosine (m^6^A) was identified as the most abundant modification in eukaryotic mRNAs^1,2^. Deposition of m^6^A occurs cotranscriptionally and is determined by the coordinated action of methyltransferases (m^6^A writers) and demethylases (m^6^A erasers)^3–5^. Their detailed mapping showed an enrichment of m^6^A near the stop codon and in the 3 ’untranslated region of the target mRNAs^2,6^.

The impact of m^6^A on mRNAs depends on their recognition by different classes of proteins called “readers”. Among the best characterized readers, the YTHDF1, 2 and 3 paralogs directly recognize m^6^A to change the translation efficiency and stability of m^6^A-containing RNAs^7,8^. Human YTHDF1, 2 and 3 are made of a C-terminal YTH-domain that binds to m^6^A separated from a N-terminal proline and glutamine-rich low-complexity domain with ∼ 70% amino acid similarity^9–11^. Despite their high level of similarity, the multiplicity of their mode of action is intensively disputed. Some studies claim their dissimilar roles, where YTHDF1 promotes mRNA translation through interactions with translational factors, YTHDF2 induces mRNA degradation, and YTHDF3 assists YTHDF1 and YTHDF2 to regulate mRNA fate^10,12,13^. By contrast, recent papers point out their redundancy with the prevailing view that YTHDF1-3 share highly similar mRNA targets and promote their degradation in a largely redundant manner^14–16^. To clarify their degree of similarity, the interactome of the YTHDF1-3 proteins has been mapped using BioID (proximity-dependent biotin identification) through their fusion with the promiscuous biotin protein ligase BirA. To date, these BioID mapping have led to the publication of contradictory observations. Zaccara et al. reported a high level of resemblance between the YTHDF interaction networks by comparing proteins enriched with C-terminal BirA fusion of YTHDF1 to those with the N-terminal BirA fusion of YTHDF2 and YTHDF3^14^. This mapping identified high-scoring interactions with mRNA silencing factors such as the CCR4-NOT deadenylase complex, the exoribonuclease XRN1, and the translational repressor DDX6^14^. By contrast, Zou et al. argued that YTHDF1, 2 and 3 have distinct partners using C-terminal BirA fusions, assuming that YTHDF1 is found with translation machineries such as initiation factors (eIFs), while YTHDF2 associates better with mRNA silencing factors, and YTHDF3 is in the margin between YTHDF1 and 2^17^.

Mechanistically, YTHDF2 remains the best-characterized m^6^A reader of the YTHDF family. Its knock-down results in the accumulation of its targets in translatable or actively translating polysome pools, pointing to a crucial role in the translational repression of its targets. Accordingly, this defect in translation is also accompanied by an increase in the global abundance of m^6^A-modified mRNAs, confirming the link existing between the number of m^6^A sites and the degradation rates of the targeted mRNA^10^. This role of YTHDF2 in the degradation of m^6^A-containing mRNAs is further supported by its capacity to recruit various mRNA silencing machineries through an effector region, namely a low-complexity segment of 385 amino acids that comprises the remainder of the protein outside of the YTH domain^10^. These silencing machineries include the CCR4-NOT complex, the RNase P/MRP endoribonuclease, and the decapping-promoting proteins UPF1/PNRC2/DCP1a^18–20^. Evidence also suggests that YTHDF2 transports its targets from the translatable pool to mRNA decay sites, such as processing bodies (P-bodies), although a recent study argues that it may only have a negligible impact in mRNA partitioning into granules^21–23^.

Here, in an attempt to decrypt the mechanism behind the multiplicity of YTHDF1-3, we present a comprehensive clarification of their interactome through an systematic BioID strategy. Notably, we report the proteomic mapping of the “effector” N-terminus of YTHDF2, which appears as a nexus for contacting both mRNA silencing factors and the translational apparatus. The spatial organization of this interactome was assessed through mapping both the N- and C-terminus of YTHDF2, and concomitantly compared with those of YTHDF1 and YTHDF3. We then describe a direct interaction between YTHDF2 and the ribosomal protein RACK1, and find that the silencing activity of YTHDF2 is selectively modulated by the codon content of its targeted mRNAs. Overall, these results have important implications for understanding the underlying multiplicity of YTHDF1-3 and could reunify apparently contradictory models.

## RESULTS

### Defining the interactome of YTHDF2

A BioID proximity mapping was first used to obtain an integrative view of the interaction network of YTHDF2. Full-length YTHDF2 (FL) was fused at its N-terminus with the abortive biotin ligase BirA and expressed as bait to capture interactors in close proximity of its N-terminus. The contribution of the effector region of YTHDF2 (AA: 1-385) was also evaluated through the generation of a truncated version of BirA-tagged YTHDF2, termed YTHDF2^ΔNT^, in which the 1-385 region was deleted. Both BirA alone and BirA-fused eGFP were selected as negative controls to monitor the non-specific background of our BioID mapping (Supplementary Figure S1A). When expressed in HEK293T cells in presence of biotin, both negative controls and BirA-fused YTHDF2 (FL and ΔNT) led to an efficient activation of biotinylation (Figure 1A). Biotinylated proteins of five independent replicates, alongside negative controls, were isolated using streptavidin-affinity purification under denaturing conditions and analyzed by quantitative mass spectrometry (MS). Our differential and statistical analysis revealed that the high-confident interactome of YTHDF2 is composed of ∼400 proteins including a high amount of factors involved in mRNA deadenylation (CCR4-NOT complex), decapping (DCP1a, EDC3), 5’ to 3’ decay (XRN1) and translational repression (GIGYF1/2, 4EHP, 4E-T, DDX6) (Figure 1B and Supplementary table 1). Numerous components of the translational apparatus, including ribosomal proteins and translation factors (eIF4G2, eIF4A, eIF3 subunits), were also detected as high-confident hits. Gene ontology (GO) analysis confirmed that both “negative regulation of translation” and “mRNA catabolic process” were among the biological processes with the most significant representation among YTHDF2 interactions (Supplementary Figure S1B)

**Figure 1.**
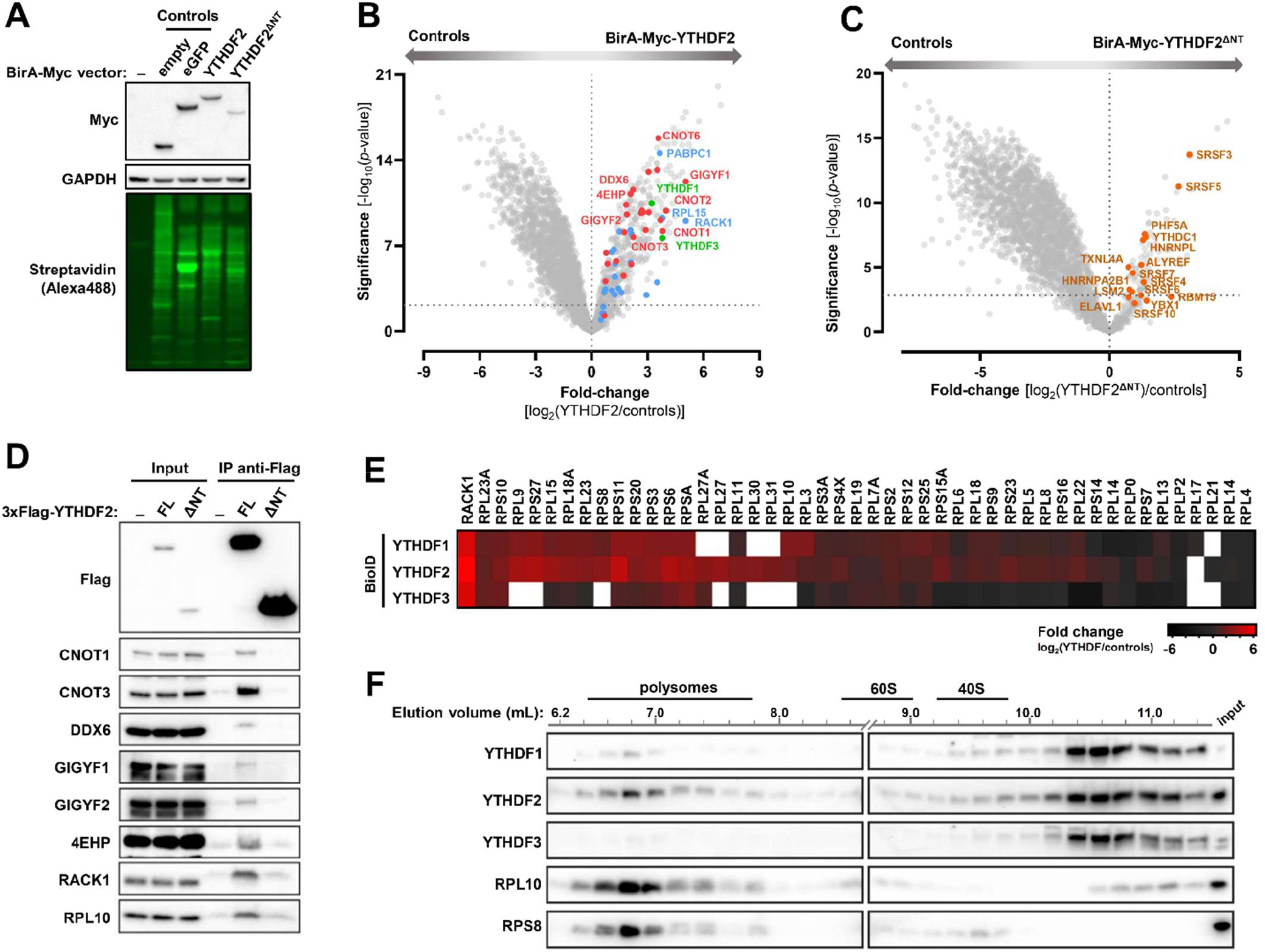
BioID mapping of the YTHDF1-3 proteins reveals their close proximity with the translation apparatus. **(A)** Western blot of lysates of non-transfected HEK293T cells (-) or transiently transfected with constructs to express either BirA-Myc alone (empty), BirA-Myc-fused eGFP, or BirA-Myc-fused YTHDF2 or a version without its N-terminus (ΔNT). Biotinylation was analysed using Alexa488-labelled streptavidin, and expression levels of the fusion proteins were analyzed with antibodies against the Myc tag. GAPDH was used as a loading control. **(B)** Volcano plot showing proteins enriched in the YTHDF2 BioID over the control BioID samples (BirA-Myc alone + BirA-Myc-fused eGFP). The logarithmic fold-changes were plotted against negative logarithmic P values of a two-sided two samples t-test. A selection of silencing factors (red) and translation factors, including ribosomal proteins (blue) is indicated. The YTHDF paralogs are shown in green. **(C)** Volcano plot showing of the interactome of a version of YTHDF2 deleted of its N-terminus (region 1-389). The plot represents the differential enrichment of proteins detected in the BirA-Myc-YTHDF2^ΔNT^ dataset (positive values) calculated against data obtained with the negative controls, BirA-Myc alone and BirA-Myc-eGFP (negative values). Proteins involved in RNA splicing are shown in orange. **(D)** Interaction of YTHDF2 with its high confidence preys. Vectors expressing 3xFlag-tagged YTHDF2, either Full-Length (FL) or ΔNT, or 3xFlag as control (-), were transfected into HEK293T cells. Extracts were immunoprecipitated with anti-Flag antibody. Total lysates (input) and IP extracts were analyzed by Western blot (WB) with the indicated antibodies. **(E)** Heatmap showing the logarithmic fold-changes of the ribosomal proteins detected in the BioID datasets of BirA-Myc-fused YTHDF1, 2 or 3 over the control BioID samples (BirA-Myc alone and BirA-Myc-fused eGFP). Note that the white color indicate that the protein was not present in the MS dataset. **(F)** Ribo Mega-SEC fractionation of a HEK293T cell lysate. Extract was fractionated using a flow rate of 0.2 ml/min on a 2,000 Å SEC column. The chromatogram is shown in Supplementary Figure S2E. Western blot was performed with the indicated antibodies.

BioID mapping of YTHDF2^ΔNT^ was performed concomitantly with its full-length version. By plotting the MS data of YTHDF2^ΔNT^ over the negative controls, we found an interactome of 106 proteins, with only 34 in common with the one of FL YTHDF2 (Figure 1C, Supplementary Figure S1C-D and Supplementary Table 2). This YTHDF2^ΔNT^dataset showed a substantial enrichment in RNA splicing and processing factors, while neither mRNA silencing factors nor ribosomal proteins were detected. To get a complementary perspective on this interaction network, BirA was also fused to the C-terminus of YTHDF2 (YTHDF2-HA-BirA). When computed over its respective negative controls (HA-BirA alone and eGFP-HA-BirA; Supplementary Figure S1E), the C-terminal BirA fusion of YTHDF2 was found to biotinylate a list of 78 proteins including 50 that were also detected with the N-terminal BirA fusion of YTHDF2 (Supplementary Figure S1F-H and Supplementary Table 3). This dataset was globally enriched in mRNA silencing factors and RNA-binding proteins such as Roquin, Pumilio and the microRNA-associated factors TNRC6A/B/C, although neither ribosomal proteins nor translation factors were detected, suggesting that the translation apparatus might be spatially too far from the BirA labelling radius at the C-terminus of YTHDF2. Overall, these data indicate that the N-terminal BirA fusion of YTHDF2 provides a broader interaction map than that of its C-terminal equivalent.

Since the deletion of the N-terminus of YTHDF2 prevents its interaction with cytoplasmic mRNA-associated machineries to favor its proximity with nuclear factors, we sought to compare the subcellular distribution of the FL and ΔNT versions of YTHDF2. In agreement with our BioID, we found that an eGFP-tagged version of YTHDF2^ΔNT^ predominantly accumulates in the nucleus of HEK293T cells, while its FL counterpart is only detected in the cytoplasm both as a diffuse signal and in punctuated foci alongside the P-body resident protein DDX6 (Supplementary Figure S1I). We then examined whether YTHDF2 utilizes its N-terminus to nucleate interactions with both mRNA silencing factors and ribosomal proteins. The stable and physical association of a Flag-tagged version of YTHDF2 with its high confident proximal interactors was examined using co-immunoprecipitation experiments (co-IP). As expected, an interaction was detected between Flag-YTHDF2 and the endogenous CNOT1 and CNOT3 subunits of the CCR4-NOT complex, the translational repressors 4EHP, GIGYF1/2 and DDX6, and the ribosomal proteins RACK1 and RPL10 (Figure 1D). Notably, these co-IPs were observed in RNase A-treated lysates, indicating that these interactions occur in an RNA-independent manner. By contrast, none of these proteins were detected following Flag-YTHDF2^ΔNT^ IP, confirming the importance of the N-terminus of YTHDF2 in mediating the interaction with both silencing factors and the ribosome.

### The ribosome is a high-confident proximal interactor of YTHDF1-3

We then sought to compare the interactome of the three YTHDF1-3 paralogs through a BioID mapping of their N-terminus (Supplementary Figure S2A). When compared to the negative controls (Myc-BirA alone and Myc-BirA-eGFP), the N-terminal BirA fusions of YTHDF1, 2 and 3 biotinylated a common set of 313 proteins showing a comparable over-representation of proteins from both mRNA silencing machineries and ribosomes (Supplementary Figure S2B-C and Supplementary Tables 4-6). In particular, global comparison of the enrichment of proteins from the ribosome among the three datasets revealed the presence of 33 ribosomal proteins with YTHDF1, 40 with YTHDF2, and 15 with YTHDF3, including 14 in common (Figure 1E). Among them, RACK1, a highly conserved factor positioned at the back side of the 40S head in the vicinity of the mRNA exit channel^24,25^, was found as the most abundant ribosomal protein shared between the YTHDF1-3 BioID (Supplementary Figure S2D).

Our BioID mapping pointed to the ribosome as a high-confident proximal interactor of YTHDF1-3. To evaluate the association of endogenous YTHDF proteins with ribosomes, HEK293T extracts were fractionated using Ribo Mega-SEC, which allows the separation of polysomes and ribosomal subunits using Size Exclusion Chromatography (SEC)^26^. The collected fractions, ranging from polysomes to smaller protein complexes, were analyzed by Western blot to visualize the fractionation profile of endogenous YTHDF1-3 along with ribosomes. Although the majority of the YTHDF1-3 pool resides in smaller complexes which did not fractionate with ribosomal components (fractions 25-31; Figure 1F and Supplementary Figure S2E), significant amounts of YTHDF2, and in a smaller extent YTHDF1, were detected in the fractions containing the ribosomal proteins (fractions 7-12). By contrast, we failed to detect YTHDF3 amongst the latter. In agreement with the BioID mapping, our co-fractionation profiles therefore indicated that the endogenous YTHDF proteins, YTHDF1 and YTHDF2 at least, can co-exist along with ribosomes.

### The ribosomal protein RACK1 directly interacts with YTHDF2

Our BioID mapping pointed out the ribosomal protein RACK1 as a high-confident interactor of the YTHDF proteins. To test whether this interaction depends on the ribosome, we generated vectors encoding Flag-tagged versions of RACK1, Wild-Type (WT) or carrying the R36D/K38E substitutions which prevent its ribosomal incorporation^27^. These vectors were expressed in HEK293T cells and RNase A-treated extracts were used for Flag IP. Following Western blot, the co-immunoprecipitation of endogenous YTHDF1 and YTHDF2 was observed with both Flag-RACK1 and its mutated version, indicating that their interaction can occur independently of the incorporation of RACK1 in the ribosome (Figure 2A). In agreement with the Ribo-Mega-SEC data (Figure 1F), we could not detect any interaction of endogenous YTHDF3 with Flag-RACK1. By contrast, CNOT1 and CNOT3, subunits of CCR4-NOT and high-confident interactors of the YTHDF proteins, were only detected with ribosomal Flag-RACK1, along with the ribosomal proteins RPS3 and RPL10, used as controls. We next sought to evaluate the contribution of RACK1 in mediating the interaction between the YTHDF proteins and the rest of the ribosome. For this purpose, RACK1-knock-out (KO) HEK293T cells (RACK1^KO^) were generated using a CRISPR/Cas9-based strategy (Supplementary Figure S3A). Vectors expressing Flag-YTHDF2, or Flag as a control, were transiently transfected in either RACK1^KO^ cells or their WT counterpart. Following Flag IP, the interaction of Flag-YTHDF2 with the ribosomal proteins RPS3 and RPL10 was more than 70% reduced in RACK1^KO^ cells in comparison with WT (Supplementary Figure S3A). A minimal co-IP remained detectable following RACK1 KO, suggesting that RACK1-independent contacts may exist between YTHDF2 and the ribosomes.

**Figure 2.**
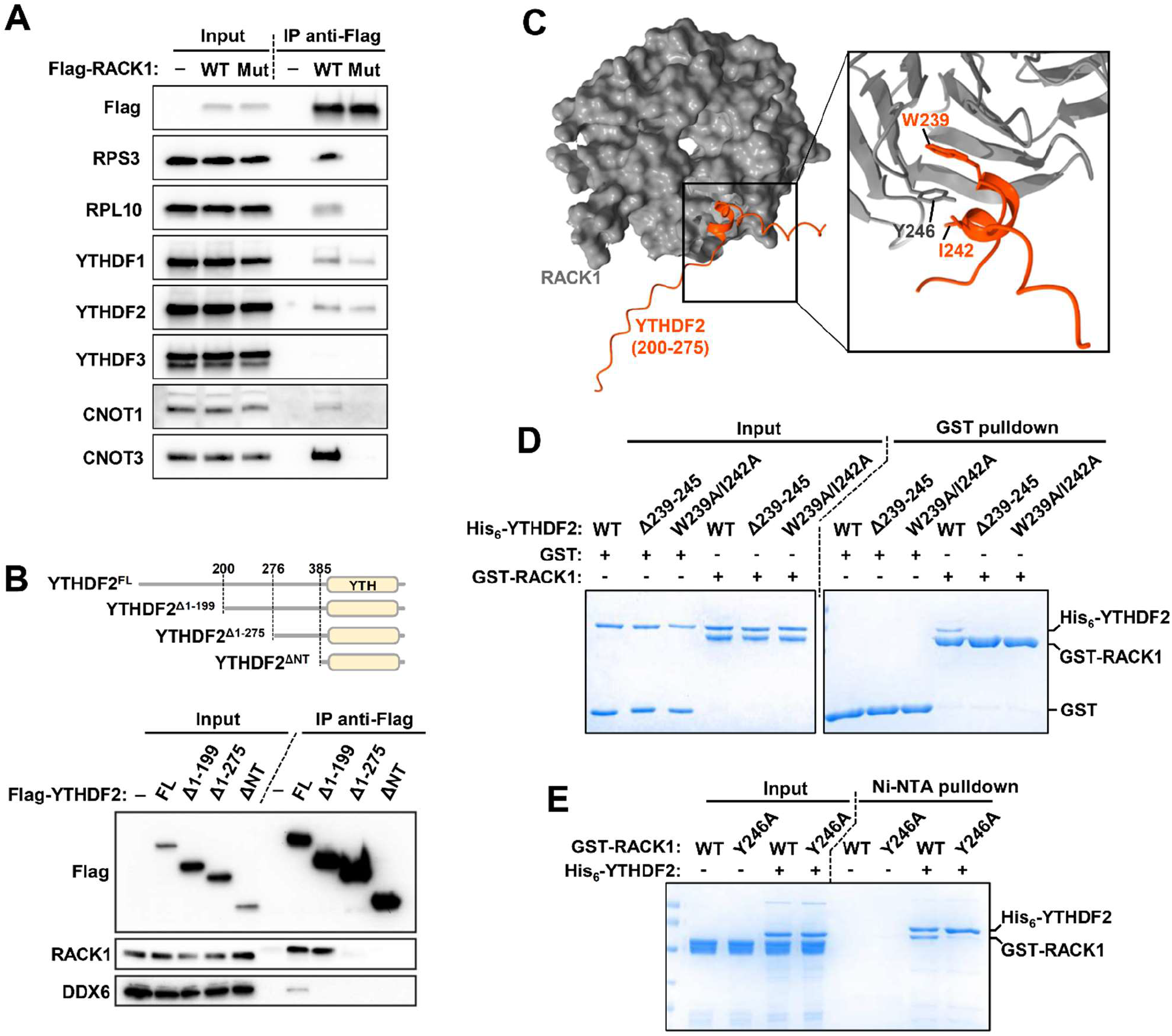
The N-terminus of YTHDF2 directly interacts with the ribosomal protein RACK1. **(A)** Western blot showing the ribosome-independent interaction between 3xFlag-RACK1 and YTHDF2. RNase A-treated extracts from cells expressing 3xFlag-RACK1, WT or carrying the R36D/K38E substitution (Mut), were used for Flag IP. Inputs and bound fractions were analyzed by Western blotting using the indicated antibodies. Empty vectors served as negative controls (-). **(B)** Top: schematic cartoon of FLAG-tagged YTHDF2 truncations immunoprecipitated in experiments shown in the bottom panel. Bottom: co-immunoprecipitation of 3xFlag-tagged versions of YTHDF2. Total lysates (input) and IP extracts were analyzed by Western blot (WB) with the indicated antibodies. **(C)** AlphaFold2-predicted structure of the 200-275 segment of YTHDF2 (orange) bound to RACK1 (gray). The insert shows an enlarged view of the putative interface involving tyrosine 246 of RACK1 (Y246) stacked between tryptophan 239 (W239) and isoleucine 242 (I242). Ribbon representations were developed using ChimeraX. **(D)** GST pull-down assay showing the interaction between recombinant GST-fused RACK1 and His_6_-YTHDF2. GST-fused RACK1 was incubated with His_6_-YTHDF2, WT or its versions harboring the double substitution W239A/I242A or deletion of the 239-245 segment (Δ239-245). GST served as negative control. The starting material (Input) and bound fractions (GST pulldown) were analyzed by SDS-PAGE followed by Coomassie blue staining. **(E)** Ni-NTA pull-down assay showing the interaction between recombinant His_6_-YTHDF2 and GST-RACK1. His_6_-YTHDF2 was incubated with GST-RACK1, WT or with the substitution Y246A. The starting material (Input) and bound fractions (Ni-NTA pull-down) were analyzed by SDS-PAGE followed by Coomassie blue staining.

We then investigated which part of YTHDF2 binds RACK1 using Flag-tagged YTHDF2 truncations and co-IP experiments. Our incremental N-terminal deletions showed that the region 200-275 of YTHDF2 is required for the maximal co-IP of endogenous RACK1 (Figure 2B). In an attempt to narrow down this minimal region, we used the AlphaFold2 structure prediction tool^28,29^. A single polypeptide comprising the RACK1 sequence fused C-terminally to the region 200-275 of YTHDF2 via a polyglycine linker was provided to AlphaFold2, and five models were generated (Supplementary Figure S3B). Remarkably, in each of the five models, YTHDF2^200–275^ was predicted as an unstructured region with the exception of a short segment (W^239^ADIAS) folding into an α-helix and positioned on the outer circumference of RACK1 (Figure 2C, Supplementary Figure S3B). RACK1 is known to adopt a seven-bladed β-propeller structure, each propeller blade consisting of a four-stranded antiparallel β-sheet^30^. In this context, the W^239^ADIAS peptide of YTHDF2 is predicted to occupy an inter-blade spacing between blades 5 and 6 with multiple contacts involving tyrosine 246 of RACK1 (Y246) stacked between tryptophan 239 (W239) and isoleucine 242 (I242) of YTHDF2 WADIAS motif. Alignment of this predicted RACK1-binding region showed that W239 and I242 are both conserved across human YTHDF1 and YTHDF3, as well as in the mouse, zebrafish and fly orthologs (Supplementary Figure S3C).

Superposition of the AlphaFold2-based model with the structure of a ribosome also showed that the YTHDF2 binding surface is accessible in ribosomal RACK1 (Supplementary Figure S3D). To validate this predicted interface, pull-down assays were performed to evaluate the interaction between recombinant Glutathione S-transferase (GST)-fused RACK1 and hexahistidine (His_6_)-YTHDF2. GST-RACK1 was incubated with His_6_-YTHDF2, WT or mutants harboring either a deletion of the 239-245 segment (Δ239-245) or the substitutions W239A/I242A. In agreement with the prediction, we observed a specific binding of WT His_6_-YTHDF2 to GST-RACK1, thus confirming their direct interaction (Figure 2D). By contrast, neither the W239A/I242A nor the Δ239-245 mutants of His_6_-YTHDF2 were bound to GST-RACK1, confirming the importance of the 239-245 region of YTHDF2 in contacting RACK1. Consistently, we also performed an Ni-NTA pull-down assay which confirmed a specific retention of WT GST-RACK1 by His_6_-YTHDF2 (Figure 2E). By contrast, the substitution of Y246 of GST-RACK1 with an alanine (Y246A) led to pronounced inhibition of this retention. Overall, these data indicate a direct interaction between the 239-245 region of YTHDF2 and an inter-blade spacing in RACK1.

### Synonymous codon usage influences YTHDF2-mediated silencing

Prior studies have reported that the degree to which mRNAs are impacted by m^6^A is dependent on their respective coding sequences in vertebrates^31^. The proximity of YTHDF2 with the ribosome prompted us to explore the possible existence of a functional connection between the mRNA coding sequence and the repressive activity of YTHDF2. The ribosome is a central determinant of mRNA stability as its translation speed directly influences the half-life of the translated mRNA (for review, see^32^). Ribosome velocity depends on the identity of the codons it translates and in particular on their optimality. To test whether YTHDF2 is involved in this process, we first interrogated how the sensitivity of mRNAs to YTHDF2 could be influenced by their codon composition using published mRNA lifetime profiling. We focused on endogenous mRNA stability profiles that have been previously published after blocking transcription in HeLa cells invalidated for YTHDF2^10^. As a metric of codon usage, the Codon Adaptation Index (CAI) of each transcript was computed and plotted based on the alteration of its mRNA lifetime following YTHDF2 knockdown. Based on this, we found that YTHDF2 preferentially induces the degradation of mRNAs with high CAI scores, while low CAI scores are enriched within the repertoire of mRNAs destabilized by the loss of YTHDF2 (Figure 3A-B), suggesting a bidirectional modulation of YTHDF2 activity based on codon usage. Early studies have shown that synonymous codon usage is correlated to GC content at third position of codons, termed GC3^33^. Accordingly, a comparable relationship was also observed when plotting GC3 levels, as well as total GC levels in the coding sequence, versus the differential mRNA lifetime following YTHDF2 knockdown (Figure 3C-D and Supplementary Figure S4A). YTHDF2-regulated mRNAs were also displayed on a scatter plot displaying the CAI of all the mRNAs detected in HeLa cells versus their GC3 score. Based on this, we confirmed the opposite distribution between the mRNAs whose lifetime was increased (high CAI, high GC3) and those having a decreased lifetime following YTHDF2 knockdown (low CAI, low GC3; Figure 3E-F). Since YTHDF2 knockdown is also known to alter translation efficiency of its targets, a similar analysis was performed using data obtained with ribosome profiling^10^. For this, we categorized the transcripts whose translation efficiency was either increased or reduced following YTHDF2 knockdown, and computed their GC3/CAI scores. In the same way as for mRNA lifetime, we found that YTHDF2 selectively favors the translation efficiency of mRNAs with low GC3/CAI scores, while it reduced that of mRNAs with high GC3/CAI scores (Supplementary Figure S4B-D). Overall, these data suggest the existence of a bidirectional activity of YTHDF2 which is correlated with the codon composition in its targeted mRNAs.

**Figure 3.**
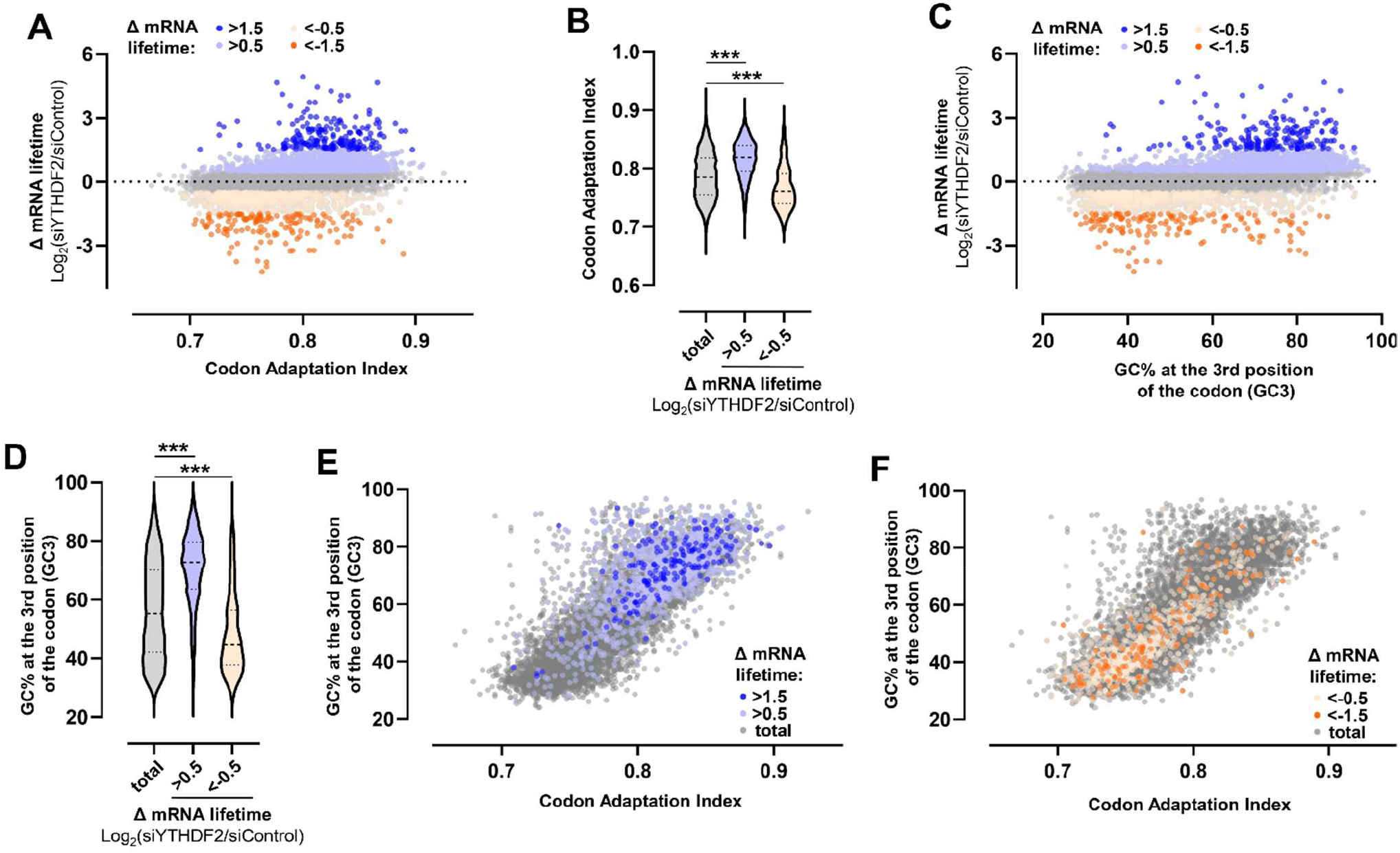
YTHDF2 sensitivity among the transcriptome is correlated with by the codon usage. **(A)** Scatter plot representing mRNA lifetime change following YTHDF2 knockdown in HeLa cells (publicly available data^10^) versus the Codon Adaptation Index of each transcript. A total of 7699 mRNAs are represented and colored according to the Δ mRNA lifetime (Log2(siYTHDF2/siControl)), including 1602 mRNAs with a Δ greater than 0.5, (197 above 1.5), and 938 with a Δ lower than -0.5 (149 below -1.5). **(B)** Violin Plot showing the CAI score for each category of mRNAs shown in (A). P value was determined by a Kolmogorov-Smirnov test: (***) P < 0.0001. **(C)** Scatter plot representing Δ mRNA lifetime following YTHDF2 knockdown (publicly available data^10^) versus the percentage of GC at the third position of the codons (GC3) of each transcript. mRNAs are colored according to the Δ mRNA lifetime (Log2(siYTHDF2/siControl)), same as in (A). **(D)** Violin Plot showing the GC3 score for each category of mRNAs shown in (C). P value was determined by a Kolmogorov-Smirnov test: (***) P < 0.0001. (**E-F**) The subsets of YTHDF2-targeted mRNAs described in (A-D) were indicated on a scatter plot showing the CAI of each mRNA detected in Hela cells versus their GC3 score (represented as grey dots). The subsets of mRNAs whose lifetime was increased (E) or decreased (F) following YTHDF2 knockdown are colored in orange and blue, respectively.

### Assessing the bidirectional codon-driven activity of YTHDF with the λN/BoxB system

To experimentally assess whether codon optimality influences the activity of the YTHDF proteins, we used the λN/BoxB system^34,35^, which is based on the artificial tethering of YTHDF onto a reporter mRNA encoding Renilla luciferase (RLuc). For this assay, YTHDF1, 2 and 3 were individually fused to the λN peptide, which has a high affinity for BoxB sequences inserted into the 3’ UTR region of the mRNA encoding luciferase (RLuc-BoxB). Co-expression of λN-YTHDFs with RLuc-BoxB mRNA induces their recruitment at the 3’ UTR, mimicking their interaction with m^6^A. To include codon usage as a parameter, we used the iCodon tool to introduce two degrees of codon optimality in the RLuc-BoxB mRNA^36^. Synonymous codon substitutions were introduced into its coding sequence to generate an optimized reporter with a high CAI score (RLuc^High^; CAI: 0.886), and a deoptimized low-score variant (RLuc^Low^; CAI: 0.733), both encoding for the same RLuc protein (Figure 4A). These reporters were individually expressed along with the λN-YTHDF proteins in HEK293T cells, together with a firefly luciferase construct (FLuc) as a transfection control (Figure 4B). As expected, we observed that the recruitment of λN-YTHDF1, 2 and 3 to the 3′ UTR of the optimized RLuc^High^ reporter markedly reduced RLuc activity compared with cells expressing only λN (∼2-fold repression, Figure 4C). Interestingly, tethering the λN-YTHDF proteins on the deoptimized RLuc^Low^ mRNA relieved their silencing activity and resulted in a ∼50% increase of RLuc production. Measurements of mRNA level by RT-qPCR showed that tethering λN-YTHDF reduced the accumulation of RLuc^High^ mRNA by ∼50%, and to a lesser extent, by only 10% that of RLuc^Low^ (Figure 4D), confirming that codon optimality selectively dictates the capacity of YTHDF1-3 to induce mRNA degradation. Besides, the YTHDF-induced upregulation in RLuc^Low^ production was thus not accompanied by a proportional increase of its mRNA level, suggesting that the translation efficiency of the non-optimal reporter mRNA was augmented by YTHDF1-3. We therefore hypothesized that this effect could be mediated by their connection with the ribosome. To test this, λN-YTHDF2 tethering was employed with the RACK1^KO^ cells in which we found that the interaction between YTHDF2 and ribosomal proteins is reduced (Supplementary figure S3A). We found that the λN-YTHDF-induced repression of the RLuc^High^ reporter remains unchanged between WT and RACK1^KO^ cells (Supplementary Figure S5A). Conversely, the upregulation of RLuc^Low^ by λN-YTHDF1, 2 and 3 was reduced by ∼50% in absence of RACK1 (Figure 4E). RT-qPCR analysis of RLuc^Low^ following λN-YTHDF2 tethering showed that this effect is not due to a significant reduction of its mRNA level in RACK1^KO^ cells compared with WT (Supplementary Figure S5B). Altogether, our tethering assays on the RLuc^High^ and RLuc^Low^ reporters indicate the existence of a selectivity in the repressive capacity of YTHDF1-3 that is directly influenced by the codon content of the targeted mRNA.

**Figure 4.**
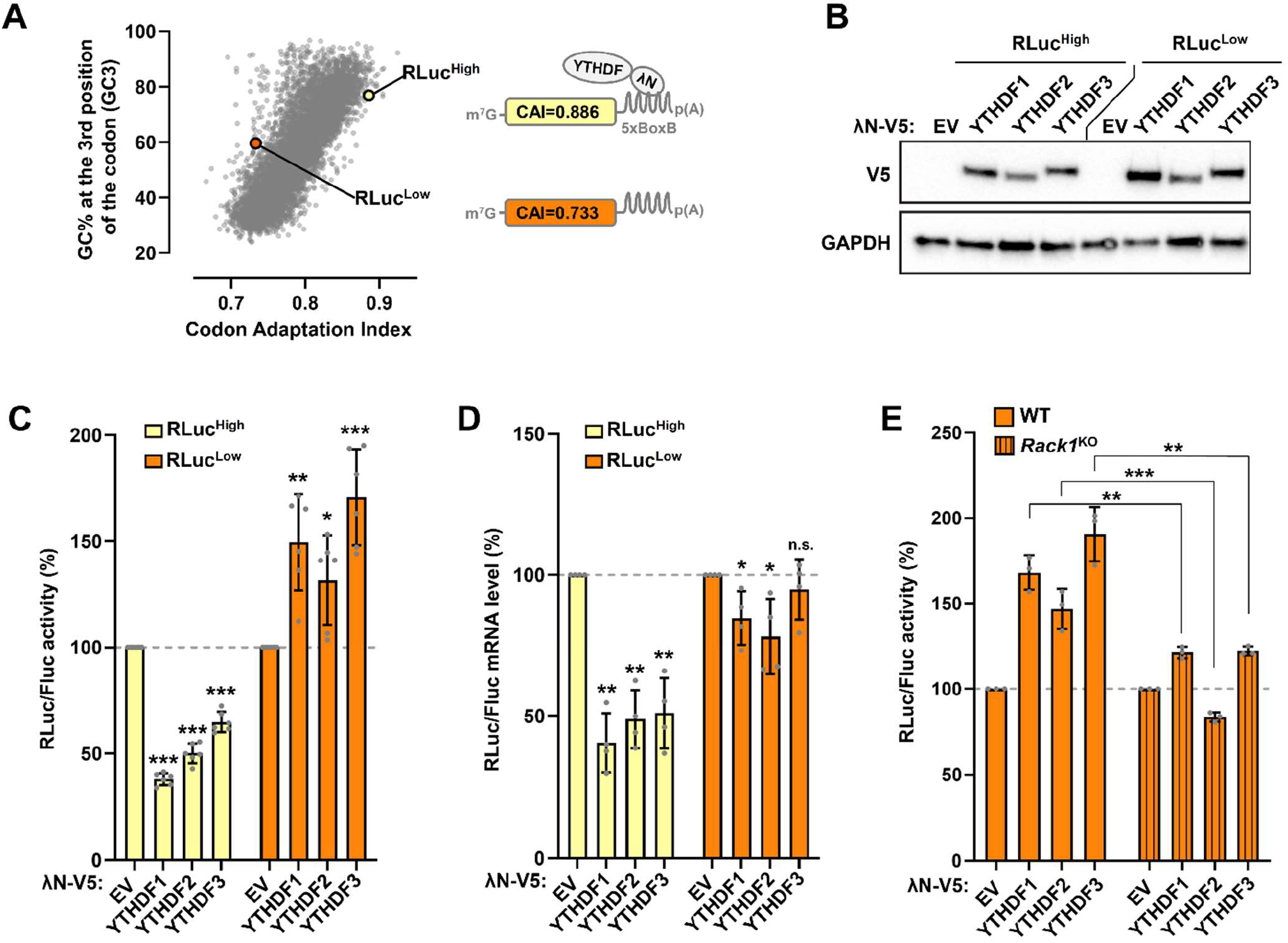
Codon usage drives YTHDF-mediated silencing of a reporter mRNA. **(A)** Schematic of the λN/BoxB tethering assay using a RLuc-5boxB reporter construct. Two degrees of codon optimality were designed for the coding region of the Renilla Luciferase (RLuc) mRNA: RLuc^High^ and RLuc^Low^ with a CAI of 0.886 and 0.733, respectively. GC3 and CAI scores of each reporter are indicated in the scatter plot representing the percentage of GC at the third position of the codon (GC3) and the CAI of the coding region for each transcript described in the UCSC database (represented as grey dots). For the tethering assay, recruitment of the YTHDF proteins to the Renilla Luciferase (RLuc) mRNA was mediated by the fused λN peptide which has a high affinity for the BoxB sequence. **(B)** Western blot showing the expression of λN-V5-fused YTHDF proteins used in the λN/BoxB tethering assay. Extracts of HEK293T cells transfected with an empty vector (EV) or plasmids encoding the λN-V5-YTHDF proteins were analyzed by Western blot with the indicated antibodies. **(C)** λN/BoxB tethering assay of the YTHDF protein on the RLuc^High^ and RLuc^Low^ reporters. HEK293T cells were co-transfected with plasmids encoding λN-V5-YTHDF1-3, RLuc^High^ or RLuc^Low^, and Firefly luciferase (FLuc). RLuc/FLuc activity values represent the mean ratio of RLuc luminescence normalized against FLuc expressed as percentage of the ratio of the cells transfected with the λN-V5 empty vector (EV). The mean values (±SD) from six independent experiments are shown and the P value was determined by two-tailed Student’s t-test: (***) P < 0.001; (**) P < 0.01; (*) P < 0.05 (compared to EV for each reporter). **(D)** Measurements by RT-qPCR to estimate the RLuc and FLuc mRNA levels from the λN/BoxB tethering assay of the YTHDF protein on the RLuc^High^ and RLuc^Low^ reporters. The mean values (±SD) from four independent experiments are shown and the P value was determined by two-tailed Student’s t-test: (**) P < 0.01; (*) P < 0.05 and n.s. means not significant (compared to EV for each reporter). **(E)** λN/BoxB tethering assay of the YTHDF protein on the RLuc^Low^ reporter in RACK1^KO^ cells. WT and *Rack1*^KO^ cells were co-transfected with indicated vectors, and RLuc/FLuc activities were analyzed as in (C).

## DISCUSSION

The action of the YTHDF1-3 proteins is a subject of continuous debate. Here, we present evidence to support a functional duality in their capacity to influence mRNA translation and stability, with an unexpected contribution of the codon content of the targeted mRNA. Based on this, we propose to derive an extended model in which the YTHDF1-3 proteins selectively induce the degradation of mRNAs enriched in optimal codons, while they promote the translation of mRNAs enriched in non-optimal codons. In this regard, our data point toward a relative redundancy of the YTHDF proteins, as we found a high degree of similarity in their interactomes and activities when tethered onto the RLuc^High^ and RLuc^Low^ reporters.

Mapping the interactomes of YTHDF1-3 revealed their unexpected proximity with the ribosomes. More that proximal, this interaction turned out to be direct through RACK1, and not brought by interactions with mRNAs, as our co-IP experiments were made with RNAse A-treated extracts. Of note, the mode of interaction between YTHDF2 and the ribosome may involve multiple interfaces, and could be more sophisticated *in cellulo* than in our *in vitro* binding assays, as an interaction between YTHDF2 and the ribosomal proteins RPS3 and RPL10 was still detectable in absence of RACK1 (Supplementary Figure S3A). Beyond the complexity of this interaction surface, our findings are consistent with previous proteomics data describing the mammalian “ribo-interactome”, namely the identification of ribosomes associated factors through affinity purification of endogenous ribosomal proteins^37^. In this report, YTHDF1/3 were listed as direct ribosome interactors, and it was proposed that they could act as anchor points on the ribosome to recruit mRNA decay factors. While it is generally assumed that mRNAs undergoing translation are protected from decay, recent work has revealed that the ribosome is a master arbiter of mRNA decay, wherein ribosome velocity serves as a major determinant of transcript half-lives^38^. Central to this emerging concept is the fact that codon usage controls ribosome traffic on mRNAs and that rare codons result in ribosome pausing^39–41^. Previous reports showed that ribosomes experiencing stalls during translation require RACK1 for proper resolution, and that RACK1 is located at the interface between collided ribosomes^42,43^. In this context, anchoring YTHDF2 onto ribosomal RACK1 could serve as a proxy to sense ribosome traffic in order to coordinate the fate of the targeted mRNA. Since slowing ribosome movement is also sensed by the CCR4-NOT complex to elicit decay of the mRNA^44^, it is tempting to speculate that the recruitment of CCR4-NOT on ribosomes, as well as its deadenylase activity, could be altered by the YTHDF2/RACK1 interaction on mRNAs enriched in non-optimal codons. Further investigation will be needed to fully resolve the interplay between the YTHDF proteins, the silencing machineries and ribosomes, and thus elucidate their impact on mRNA fate.

In agreement with previous studies^45^, our reanalysis of published datasets indicated that YTHDF2 preferentially induces the degradation of GC-rich mRNAs at transcriptome-wide level. Conversely, a striking feature of the mRNAs that tend to be less stable, as well as less translated, following YTHDF2 knock-down is that they correspond to an AU-rich subset of the transcriptome. Based on our tethering assay, this codon-driven selectivity of YTHDF2 can be extrapolated to YTHDF1 and YTHDF3, and turned out to be RACK1-dependent in the case of the AU-rich RLuc^Low^ reporter. Beyond its impact in mRNA translation and stability, AU content is known to target mRNA for P-body localization. Notably, P-bodies accumulate mostly AU-rich mRNAs with a specific codon usage associated with a low protein yield^45,46^. The YTHDF proteins can also promote m^6^A-modified mRNAs entering P-bodies. Central to this phenomenon is the propensity of the YTHDF proteins to undergo liquid-liquid phase separation (LLPS) through a disordered proline-glutamine-rich region in their effector domain (AA: 230−383 in YTHDF2)^21,22,47^. Since the ribosome is not detected in P-bodies^46^, it is plausible that the RACK1-dependent activation of AU-rich mRNAs by YTHDF2 occurs outside P-bodies. The RACK1-binding region of YTHDF2 (AA: 239-245) is located within its LLPS-inducing segment, and it is therefore tempting to speculate that the YTHDF2/RACK1 interaction could reduce condensate assembly around the methylated mRNAs in order to isolate it from silencing granules and to maintain it in the translatable pool of transcripts. Additional experiments will be mandatory to test if RACK1 affects the partitioning of AU-rich targets of YTHDF2 out of P-bodies. Alternatively, further investigation will be needed to verify whether YTHDF2 and RACK1 could co-exist in phase separated condensates. Recent studies found that granules can positively impact co-translation events, and active translation can occur in granules^48–51^. These findings may open the door to understanding how a physical intimacy between YTHDF action and translation apparatus shapes the fate of mRNAs.

## AUTHORS CONTRIBUTIONS

C.M., L.Z. N.P. and Cl.C. performed the experiments. L.Z., M.G., N.P. and Ch.C. and Cl.C. designed research and analyzed the experiments, G.B. helped with Ribo Mega-SEC experiments, and C.M., L.Z., and Cl.C. wrote the paper.

## ACKNOWLEDGMENTS

We thank N. Ulryck for helpful technical assistance. W. Filipowicz (FMI, Basel) and N. Gehring (University of Cologne) are acknowledged for their gifts of pCI-RL (5BoxB) and pCI-λN-V5, respectively. The pET-28c(+)-YTHDF2 vector was a gift from Samie Jaffrey (Addgene plasmid 102275), pcDNA3.1 mycBioID from Kyle Roux (Addgene plasmid 35700), pSpCas9(BB)-2A-Puro from F. Zhang (Addgene plasmid 62988). Cl.C. acknowledges financial supports from Les Entreprises contre le cancer – Gefluc – Paris Île-de-France, Fondation ARC (PJA2 2020) and Agence Nationale pour la Recherche [grant ANR-22-CE12-0029]. L.Z. is supported by a PhD fellowship from the China Scholarship Council (CSC). M.G. and Cl.C. acknowledge financial supports from Ecole Polytechnique, the Centre National pour la Recherche Scientifique, the Agence Nationale pour la Recherche [grant ANR-16-CE11-0003]. BioID and MS experiments were also supported by Agence Nationale pour la Recherche via the French Proteomic Infrastructure ProFI FR2048 [grant ANR-10-INSB-08-03]. The imaging facility of Laboratoire d’Optique et Biosciences (Ecole Polytechnique) was partly supported by Agence Nationale de la Recherche [ANR-11-EQPX-0029 Morphoscope2].

## Materials availability

All newly created cell populations generated in this study are available upon request.

## Data availability

All data reported in this paper and any additional information required to reanalyze the data will be shared by the lead contact upon request.

## METHODS

### Cell culture

HEK293T cells (Sigma-Aldrich) were routinely maintained in DMEM supplemented with 10% FBS and 2% penicillin/streptomycin in a humidified atmosphere of 5% CO2 at 37 °C.

### Plasmid cloning

For fusion to the C-terminus of BirA(R118G), the human YTHDF1, 2, 3 ORFs provided by Mission TRC3 Human LentiORF Clones TRCN0000477232, TRCN0000468616, TRCN0000473510, respectively, were cloned by PCR as EcoRI–BamHI fragments into the vector pcDNA3.1 mycBioID^52^ (Addgene plasmid #35700) in frame with BirA. Similarly, to generate the BirA-fused eGFP used as a negative control for BioID, a fragment containing the eGFP coding sequence was obtained by PCR from the pEGFPC1 and inserted at the EcoRI–BamHI sites of the pcDNA3.1 mycBioID vector. pCI-eGFP-YTHDF2 was generated by inserting the eGFP sequence at the NheI-XhoI site of the pCI-Neo vector (Promega) and then by adding a fragment encoding YTHDF2 at the XhoI-NotI site. To create 3xFlag expression plasmids, an annealed oligonucleotide encoding the triple Flag peptide was inserted as a NheI–XhoI fragment into the pCI-Neo plasmid. YTHDF1, 2 and 3 sequences were then inserted in frame with the triple Flag sequence at the XhoI-NotI sites. PCR amplifications were also used to create truncated fragments of YTHDF2 which were inserted in the pCI-3xFlag vector. For tethering assays, fragments containing the YTHDF1, 2 or 3 sequence were obtained by PCR and inserted into the pCI-λNV5 at the XhoI-NotI sites. As previously described^35^, a pCI-RLuc-BoxB vector containing RLuc^High^ (CAI: 0.886) was used. The deoptimized low-score RLuc variant (RLuc^Low^; CAI: 0.733) was synthetized as a gBlock gene fragment (Integrated DNA Technologies) and inserted at the NheI-XbaI of pCI-RLuc-BoxB instead of the RLuc^High^ fragment. For bacterial expression of His-tagged YTHDF2, the pET-28c(+)-YTHDF2 vector (Addgene plasmid #102275) was used^53^. For LLPS assays, a PCR fragment encoding a monomeric eGFP (A207K) was inserted at the NdeI site of pET-28c(+)-YTHDF2. The full-length cDNA encompassing the coding region of human RACK1 was obtained by RT-PCR using total RNA from HEK293T cells and inserted at the XhoI-NotI sites of the pCI-Neo-3xFlag vector, and at the SalI-NotI of the pGEX-6P-1 vector (Amersham) in frame with the GST coding sequence. For the bacterial expression of GST-fused mCherry2-RACK1, a gBlock gene fragment (Integrated DNA Technologies) encoding the mCherry2-RACK1 fusion protein was also inserted at the SalI-NotI of the pGEX-6P-1. Amino acid substitutions and deletions were introduced by site-directed mutagenesis using the QuikChange Kit (Agilent). Sequences of the primers used are listed in the Supplementary Table S7.

### BioID and affinity purification

BioID experiments were performed as previously described with minor changes^54^. Briefly, HEK293T cells were grown in 15 cm dishes and transfected with 15 µg of plasmids expressing the BirA-fused baits. The day after, biotinylation was induced by adding biotin (50 µM) in the medium. After 24 h of treatment, cells were rinsed once on the plate with 20 mL of PBS, then scraped into 1 mL of PBS. Cell pellets were collected by centrifugation (500 × g for 5 min) and resuspended in 1 mL of ice-cold lysis buffer (50 mM Tris⋅HCl, pH 7.5, 150 mM NaCl, 1% IGEPAL® CA-630, 0.4% SDS, 1.5 mM MgCl2, 1 mM EGTA, benzonase, and cOmplete EDTA-free Protease inhibitor). Cells were dispersed with a P1000 pipette tip (∼10–15 aspirations), lysates were rotated at 4 °C for 30 min and then centrifuged at 16,000 × g for 20 min at 4 °C. Supernatant was collected into new tubes for affinity purification. Samples were incubated with 20 µL (packed beads) of streptavidin-Sepharose (GE) (equilibrated in lysis buffer) with rotation overnight at 4 °C. Beads were collected (500 × g for 2 min), the supernatant was discarded, and the beads were transferred to new tubes in 500 µL of lysis buffer. Beads were washed once with SDS wash buffer (50 mM Tris⋅HCl, pH 7.5, and 2% SDS), twice with lysis buffer, and three times with 50 mM ammonium bicarbonate pH 8.0 (all wash volumes: 500 µL with centrifugations at 500 × g for 30 s).

### Proteomic sample preparation

Affinity purified proteins retained on beads were resuspended in 100 µL of NH_4_HCO_3_ at 25 mM containing 2 µg of Trypsin/Lys-C mix (Mass Spec Grade, Promega, Madison, WI, USA). Digestion was performed under agitation at 37°C during 4h. Then, the same enzyme amount was added to the solution and the second digestion step was conducted overnight at 37°C under agitation. Supernatant was collected after a centrifugation at 500 × g for 2 min, and beads were washed with water. Both solutions were pooled. The enzymatic digestion was stopped by adding 6 µL of pure formic acid (2% final concentration). Samples were then centrifuged at 16,000 × g for 5 min and supernatants were collected (90% of the solution). Finally, the samples were vacuum dried, and then resolubilized in 400 µL of H_2_O/acetonitrile/formic acid (98/2/0.1 v/v/v).

### NanoLC-MS/MS Analysis

NanoLC-MS/MS analyses were performed on a nanoAcquity UPLC device (Waters Corporation, Milford, USA) coupled to a Q-Exactive HF-X mass spectrometer (Thermo Fisher Scientific, Bremen, Germany). Peptides were loaded on a symmetry C18 pre-column (20 mm × 180 μm with 5 μm diameter particles, Waters) before being separated on an ACQUITY UPLC BEH130 C18 column (250 mm × 75 μm with 1.7 μm diameter particles). The solvent system consisted of 0.1% formic acid in water (solvent A) and 0.1% formic acid in acetonitrile (solvent B). The samples were loaded into the enrichment column over 3 min at 5 μL/min with 99% of solvent A. Peptides were then eluted at 450 nL/min with the following gradient of solvent B: from 1 to 8% over 2 min, 8 to 35% over 77 min, and 35 to 90% over 1 min. The MS capillary voltage was set to 2 kV at 250 °C. The system was operated in a data-dependent acquisition (DDA) mode with automatic switching between MS (mass range 375-1500 m/z with R = 120,000 at 200 m/z, automatic gain control fixed at 3 × 10^6^ ions, and a maximum injection time set at 60 ms) and MS/MS (mass range 200-2000 m/z with R = 15,000 at 200 m/z, automatic gain control fixed at 1 × 10^5^, and the maximal injection time set to 60 ms) modes. The twenty most abundant peptides were selected on each MS spectrum for further isolation and higher energy collision dissociation fragmentation (normalized collision energy set to 27), excluding unassigned, singly charged and over seven times charged ions. The dynamic exclusion time was set to 40 sec, and peptide match selection was turned on preferred.

### Quantitative proteomic data analysis

Raw data collected during nanoLC-MS/MS were processed using MaxQuant software (version 1.6.6.0)^55^. Peaks were assigned with the Andromeda search engine with trypsin specificity. The database used for the searches was extracted from UniProtKB-SwissProt and included all *Homo sapiens (Human)* entries (22 July 2019; Taxonomy ID=9606; 20,409 entries). The minimum peptide length required was seven amino acids and a maximum of one missed cleavage was allowed. The precursor mass tolerance was set to 20 ppm for the first search and 4.5 ppm for the main search. The fragment ion mass tolerance was set to 20 ppm. Methionine oxidation and acetyl (Protein N-term) were set as variable modifications. The maximum false discovery rate was 1% for peptides and proteins with the use of a decoy strategy. The “match between runs” option was desactivated. Unique peptides were used but modified peptides as well as their unmodified counterparts, were excluded from protein quantification. We used the “proteingroups.txt” file with intensities (non-normalized intensities). The dataset was deposited to the ProteomeXchange Consortium via the PRIDE partner repository with the dataset identifier PXD046957.

### Statistical data analysis of proteomic data

Statistical analyses were performed using ProStaR software^56^. Proteins identified in the reverse and contaminant databases and those only identified by site were removed. Moreover, only proteins for which 5 intensity values were available in a single condition were kept. After log2 transformation, intensities were normalized within condition using vsn method and imputation of missing values were performed. For each sample, the slsa algorithm was used for the POV (Partially Observed Values) imputation, and missing values were replaced by the 2.5 percentile value for the MEC (Missing on the Entire Condition); statistical testing was performed using Limma. Benjamini-Hochberg method was used to adjust p-values for multiple testing and differentially expressed proteins were sorted out using a p-value threshold that guarantees a FDR below 1%.

### Western blot and antibodies

Proteins were separated by SDS-PAGE and transferred onto low-fluorescence PVDF membranes. The membranes were blocked in PBS containing 5 % non-fat milk and 0.1% Tween 20 for 30 min at room temperature. Blots were probed with the following antibodies: rabbit anti-eIF4E2 (4EHP) (Proteintech, catalog no. 12227-1-AP), rabbit anti-YTHDF1 (Proteintech, catalog no. 17479-1-AP), rabbit anti-YTHDF2 (Proteintech, catalog no. 24744-1-AP), rabbit anti-YTHDF3 (Proteintech, catalog no. 25537-1-AP), rabbit anti-eIF4ENIF1 (4E-T; Abcam, catalog no. ab55881), rabbit anti-DDX6 (Proteintech, catalog no. 14632-1-AP), rabbit anti-CNOT1 (Proteintech, catalog no. 14276-1-AP), mouse anti-Flag (Sigma, catalog no. F1804), rabbit anti-CNOT3 (Proteintech, catalog no. 11135-1-AP), rabbit anti-GIGYF1 (Bethyl, catalog no. A304-133A), rabbit anti-GIGYF2 (Proteintech, catalog no. 24790-1-AP), mouse anti-GAPDH (Proteintech, catalog no. 60004-1-Ig), rabbit anti-XRN1 (Proteintech, catalog no. 23108-1-AP), rabbit anti-RACK1 (Cell Signaling Technology, catalog no.5432), rabbit anti-RPL10 (Cell Signaling Technology, catalog no. 72912), rabbit anti-RPS3 (Proteintech, catalog no. 11990-1-AP), anti-V5 tag (Invitrogen, catalog no.R960-25).

### Extract Preparation and Immunoprecipitation

Cells were resuspended in a lysis buffer containing 20 mM HEPES-KOH, pH 7.5, 100 mM NaCl, 2.5 mM MgCl2, 0.5% IGEPAL® CA-630, 0.25% sodium deoxycholate, supplemented with cOmplete EDTA-free Protease inhibitor and phosphatase inhibitor Cocktail (Roche), and incubated for 20 min on ice. The lysate was clarified by centrifugation at 15,000 × g for 10 min at 4 °C. One milligram of extract was used for immunoprecipitation with the indicated antibodies. Thirty microliters of pre-equilibrated Anti-FLAG® M2 Magnetic Beads (Millipore, catalog no. M8823) and RNase A (Thermo Scientific) were added, and the mixtures were rotated overnight at 4 °C. Beads were washed five times with lysis buffer and directly resuspended in protein sample buffer for Western blot analysis.

### CRISPR/cas9-mediated genome editing

CRISPR-Cas9-mediated genome editing of HEK293 cells was performed as previously described^57^. The following oligonucleotides encoding a small guide RNA cognate to the coding region of *Rack1* gene were used: 5’-CACCGTGTCAACCGCACGTCTATGC and 5’-AAACGCATAGACGTGCGGTTGACAC. These oligos which contain BbsI restriction sites were annealed creating overhangs for cloning of the guide sequence oligos into pSpCas9(BB)-2A-Puro (PX459) V2.0 (Addgene plasmid #62988) by BbsI digestion. To generate KO HEK293T cells, we transfected 700,000 cells with the pSpCas9(BB)-2A-Puro plasmid. 24 hr after transfection, puromycin was added in the cell medium to 1.5 µg/mL final concentration. After 72 h, puromycin-resistant cells were isolated into 96-well plates to obtain monoclonal colonies. Clonal cell populations were analyzed by WB for protein depletion.

### Polysome fractionation by Ribo Mega-SEC

Ribo Mega-SEC was performed as previously described with minor changes^26^. Cells in two 15 cm dishes (70% confluency) were treated with 100 μg/ml cycloheximide for 5 min, washed with ice-cold PBS containing 50 μg/ml cycloheximide, scraped on ice, collected by centrifugation, lysed in 600 μl of polysome extraction buffer (20 mM Hepes-NaOH (pH 7.4), 130 mM NaCl, 10 mM MgCl_2_, 1% CHAPS, 2.5 mM DTT, 50 μg/ml cycloheximide, 20 U RNaseIn RNase inhibitor, cOmplete EDTA-free Protease inhibitor), incubated for 15 min on ice, and centrifuged at 17,000 *g* for 10 min (all centrifugations at 4°C). Supernatants were filtered through 0.45 μm Ultrafree-MC HV centrifugal filter units by 12,000 *g* for 2 min before injecting to SEC. For SEC, an Agilent Bio SEC-5 2,000 Å column (7.8 × 300 mm with 5 μm particles) was equilibrated with two column volumes of filtered SEC buffer (20 mM Hepes-NaOH (pH 7.4), 60 mM NaCl, 10 mM MgCl_2_, 0.3% CHAPS, 2.5 mM DTT) and 100 μl of 10 mg/ ml of filtered bovine serum albumin (BSA) solution diluted by PBS was injected once to block the sites for non-specific interactions. 500 µl of cell lysates were injected onto the column. The flow rate was 0.2 ml/min and 48 × 100 μl fractions were collected in a low-protein binding 96-deep-well plate (Eppendorf) at 4°C. The chromatogram was monitored by measuring UV absorbance at 215, 260 and 280 nm. 20 µl of each fraction were used for western blotting.

### Structure prediction using AlphaFold2

The predicted structure of the YTHDF2/RACK1 interaction was performed using AlphaFold2 as implemented in ColabFold^28,29^. A chimeric sequence in which full-length RACK1 was concatenated at the C-terminal end to YTHDF2 (AA: 200-275) with a 30-glycine linker, and used as input in default settings and in absence of any imposed constraints to generate five models.

### Codon usage analysis

BioMART was used to retrieve the coding sequences of 19,068 human transcripts (GRCh38.p14) from *Ensembl*. The codon adaptation index and GC level were calculated using the CAIcal tool (genomes.urv.es/CAIcal/). Statistical analysis and plots were obtained with GraphPad Prism software.

### Confocal microscopy scanning

HEK293T cells were grown in U-Slide ibiTreat coverslip multi-well chambers and transfected with 20 ng of pCI-eGFP-YTHDF2 (FL or ΔNT) vector. After 24 h, cells were fixed with 4% paraformaldehyde and permeabilized in PBS containing 0.1% Triton X100, blocked in blocking buffer (1% BSA, 22.5 mg/mL glycine in PBS + 0.1% Tween 20) for 30 min. Anti-DDX6 antibody was added onto the cells in blocking buffer for 1 h at 37 °C. After three washes in PBS, cells were incubated with secondary antibody coupled with Alexa Fluor 568 (Thermo Fisher Scientific, catalog no. A-11011) for 45 min at 37°C, and mounted on glass slides in mounting solution containing DAPI (Fluoroshield, Sigma-Aldrich). Fluorescence was visualized under a LEICA-SP8ST-WS confocal microscope.

### Expression and purification of His_6_-YTHDF2

His_6_-YTHDF2, FL or its mutants (W239A/I242A and Δ239-245) were expressed in *E. coli* BL21 (DE3) Codon+ (Agilent). Large-scale expression was done in 1 L of auto-inducible terrific broth media (ForMedium AIMTB0260) supplemented with kanamycin (50 μg/mL) and chloramphenicol (25 μg/mL), first at 37°C for 3 h and then at 18°C overnight. Cells were collected, pelleted and then resuspended in 30 mL of lysis buffer (25 mM HEPES pH 7.5, 300 mM NaCl and 20 mM Imidazole) supplemented with one protease inhibitor tablet (Roche), 0.5 mM PMSF and 30 μL benzonase nuclease (Millipore Sigma). The cells were lysed by sonication on ice and the lysate clearance was performed by centrifugation at 20,000 g for 30 min. The supernatant was applied on Talon Metal Affinity Resin (Clontech) pre-equilibrated with the lysis buffer and incubated at 4°C on a rotating wheel for 1 h, followed by a washing step with 30 mL of washing buffer (25 mM HEPES pH 7.5 and 1 M NaCl). His6-YTHDF2 was eluted by the addition of 15 mL of elution buffer (25mM HEPES pH 7.5, 300 mM NaCl and 400 mM Imidazole), followed by concentrating up to 10 mL by a 30 kDa cutoff concentrator. The sample was then diluted to 50 mL using Hitrap S buffer A (50 mM MES pH 6.0 and 50mM NaCl), followed by loading on a 5 mL an HiTrap S FF column (Cytiva) and eluted using a NaCl linear gradient from 50 mM (100% Hitrap S buffer A) to 1 M (100% Hitrap S buffer B: 50 mM MES pH 6.0, and 1 M NaCl). The fractions containing His_6_-YTHDF2 protein were collected, concentrated up to 5 mL, and injected on a Superdex 200 increase 10/300 size-exclusion column (Cytiva) with Gel filtration buffer (25 mM HEPES pH 7.5, 300 mM NaCl and 5 mM β-mercaptoethanol). The fractions containing His_6_-YTHDF2 were collected and concentrated.

### Expression and purification of GST-RACK1

Expression of GST-RACK1 and its mutant Y246A was carried out in *E. coli* BL21 (DE3) Codon+ (Agilent) in 1 L of auto-inducible terrific broth media (ForMedium AIMTB0260) supplemented with ampicillin at 100 μg/mL and chloramphenicol at 25 μg/mL. Cultures were incubated at 18°C for 20 h. Bacteria were harvested by centrifugation and resuspended in lysis buffer (25 mM HEPES pH 7.5, 50 mM NaCl, 5 mM β-mercaptoethanol). Cell lysis was performed by sonication on ice. After centrifugation for 30 min at 20,000 g at 4°C, clarified samples were transferred to batch-bind with Glutathione SepharoseTM 4B (Cytiva) resin for 1 h at 4°C followed by a washing step with 30 mL of washing buffer (25 mM HEPES pH 7.5, 50mM NaCl and 5 mM β-mercaptoethanol) supplemented with 10 mM ATP. GST-RACK1 was eluted with lysis buffer supplemented with 20 mM GSH (pH 7.0) and further purified on an HiTrap Q FF column (Cytiva) using a linear gradient of 100% Hitrap buffer A (20 mM Tris/HCl pH 8.0, 50 mM NaCl and 5 mM β-mercaptoethanol) to 100% Hitrap buffer B (20 mM Tris/HCl pH 8.0, 1 M NaCl and 5 mM β-mercaptoethanol). The peak fractions corresponding to GST-RACK1 were pooled, concentrated up to 5 mL, and injected on a Superdex 75 increase 10/300 size-exclusion column (Cytiva) with Gel filtration buffer (20 mM Tris/HCl pH 8.0, 200 mM NaCl and 5 mM β-mercaptoethanol). The fractions containing GST-RACK1 were concentrated and used for pull-down assays.

### Pull-down assays

Both Ni-NTA and GST pull-down experiments were performed by incubating 1 nmol of His_6_-YTHDF2, FL or mutants, with the same molar amount of GST-RACK1. GST was used as negative control. All proteins were free of nucleic acids according to the OD 280 nm /OD 260 nm ratio. For Ni–NTA pull-down, binding buffer (25 mM HEPES pH 7.5, 150 mM NaCl and 50 mM imidazole) was added to a final volume of 60 μL. The reaction mixtures were incubated on ice for 1 h. After that, 10 μL was withdrawn and used as an input fraction for SDS-PAGE analysis. The remaining 50 μL were incubated at 4°C for 2 h with 40 μg of pre-equilibrated HisPur Ni–NTA magnetic beads (Thermo Scientific) in a final volume of 200 μL. Beads were then washed three times with 500 μL of binding buffer. Bound proteins were eluted with 50 μL of elution buffer (25 mM HEPES pH 7.5, 150 mM NaCl, 250 mM imidazole and 10% glycerol). Samples were resolved on SDS-PAGE and visualized by Coomassie blue staining. For GST pull-down, the supernatant was applied on pre-equilibrated Glutathione SepharoseTM 4B resin (Cytiva) and incubated at 4°C on a rotating wheel for 1 h, followed by a washing step with 1 mL of lysis buffer (50 mM HEPES pH 7.5, 150mM NaCl and 10% Glycerol). Retained proteins were eluted by 50 μL of elution buffer (50 mM HEPES pH 7.5, 150mM NaCl, 10% Glycerol and 20mM GSH), followed by SDS-PAGE and Coomassie blue staining.

### Tethering and luciferase assays

HEK293T cells were transfected with 20 ng of RLuc-5BoxB, 5 ng of FLuc, and 100 ng of λN-fusion constructs per well in a 24-well plate by using Lipofectamine 2000 (Thermo Scientific, 11668019) according to the manufacturer’s instructions. Cells were lysed 24 h after transfection and luciferase activities were measured with the Dual-Luciferase Reporter Assay System (Promega) in a GloMax 20/20 luminometer (Promega). RLuc/FLuc activity values represent the mean ratio of RLuc luminescence normalized against FLuc expressed as percentage of the ratio of the cells transfected with the λN-V5 empty vector.

