## Supplementary information for "The YTHDF1-3 proteins are bidirectionally influenced by the codon content of their mRNA targets"

**Supplementary Figure S1 –** BioID mapping of the interactome of YTHDF2

**Supplementary Figure S2 –** BioID mapping of the YTHDF1-3 paralogs shows their proximity with ribosomes

**Supplementary Figure S3 –** RACK1 partially mediates the interaction between YTHDF2 and ribosomes

**Supplementary Figure S4 –** YTHDF2 sensitivity among the transcriptome correlates with the GC content

**Supplementary Figure S5 –** Tethering the YTHDF proteins in RACK1$^{KO}$ cells

**Supplementary Table S7 –** Oligonucleotides used in this study

### Supplementary Figure S1

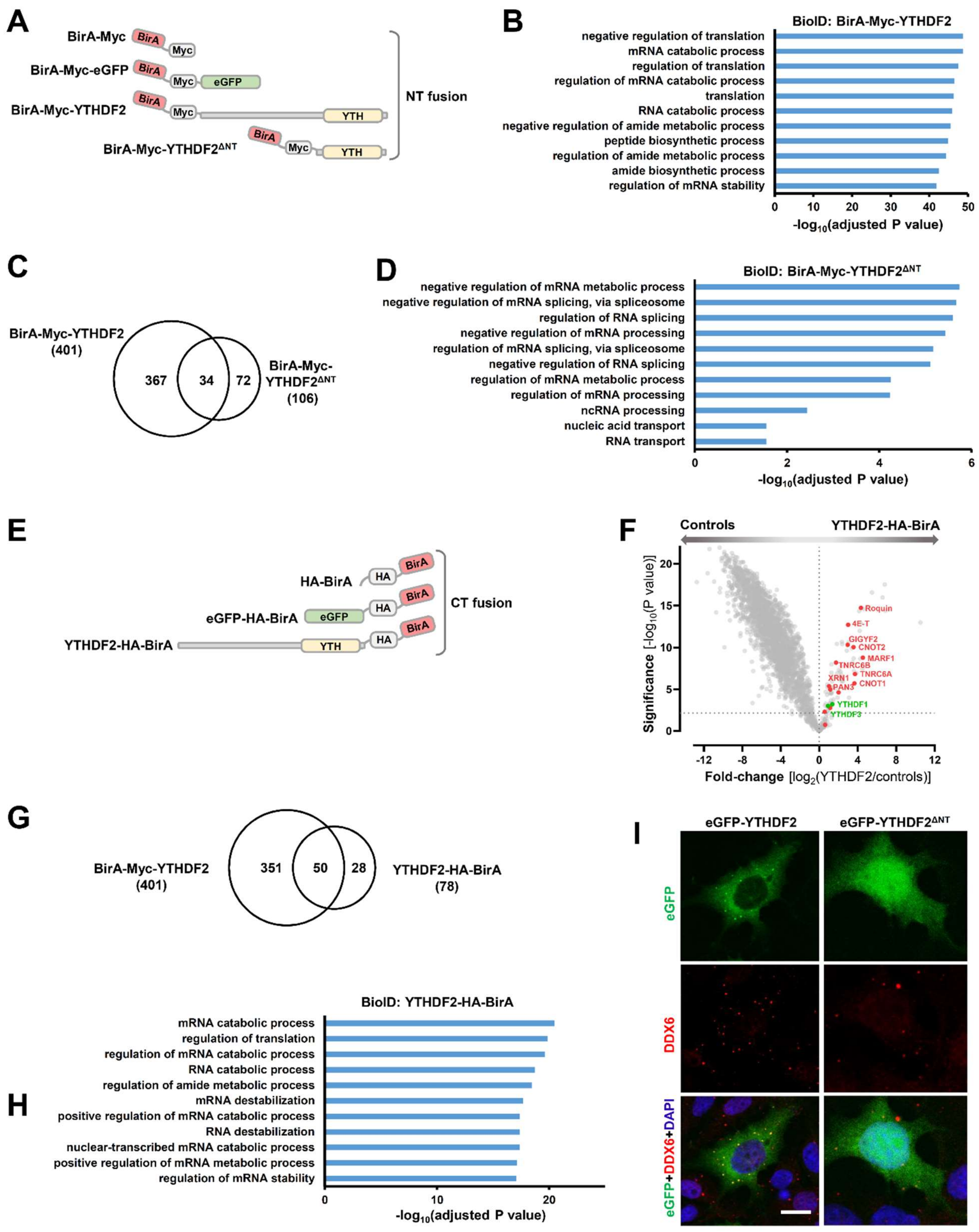


**Supplementary Figure S1 – BioID mapping of the interactome of YTHDF2.** (**A**) Schematic cartoon of the BirA-Myc fused constructs used to generate the BioID datasets presented in Figure 1A-C and panels B-D hereafter. Both BirA and the Myc tag were fused at the N-terminus (NT) of eGFP, YTHDF2 and YTHDF2$^{\Delta NT}$.

(**B**) GO analysis of the BirA-Myc-YTHDF2 proximal proteins, related to data presented in Figure 1B and Supplementary Table 1. The most significantly enriched biological processes identified by g:Profiler software are presented. (**C**) Venn diagram comparing the BioID datasets of BirA-Myc-YTHDF2 and BirA-Myc-YTHDF2$^{\Delta NT}$. (**D**) Graph showing GO analysis of the BirA-Myc-YTHDF2$^{\Delta NT}$ proximal proteins with the most significantly enriched biological processes, mostly related to splicing. (**E**) Depiction of the BirA-HA fused constructs used to generate the BioID datasets presented in the panels F-H hereafter. Both BirA and the HA tag were fused at the C-terminus (CT) of eGFP and YTHDF2. (**F**) Volcano plot showing proteins enriched in the YTHDF2-HA-BirA BioID (CT fusion) over the control BioID samples (HA-BirA alone + eGFP-HA-eGFP). The logarithmic fold-changes were plotted against negative logarithmic P values of a two-sided two samples t-test. A selection of silencing factors (red) is indicated. The YTHDF paralogs are shown in green. (**G**) Venn diagram comparing the BioID datasets of BirA-Myc-YTHDF2 (NT fusion) and YTHDF2-HA-BirA (CT fusion). (**H**) GO analysis of the YTHDF2-HA-BirA proximal proteins showing the most significantly enriched biological processes. (**I**) Confocal analysis of HEK293T cells expressing eGFP-fused YTHDF2 or YTHDF2$^{\Delta NT}$, and immunostained for DDX6 (red). Nuclei were stained with DAPI (blue). Scale bar = 5µm.

### Supplementary Figure S2

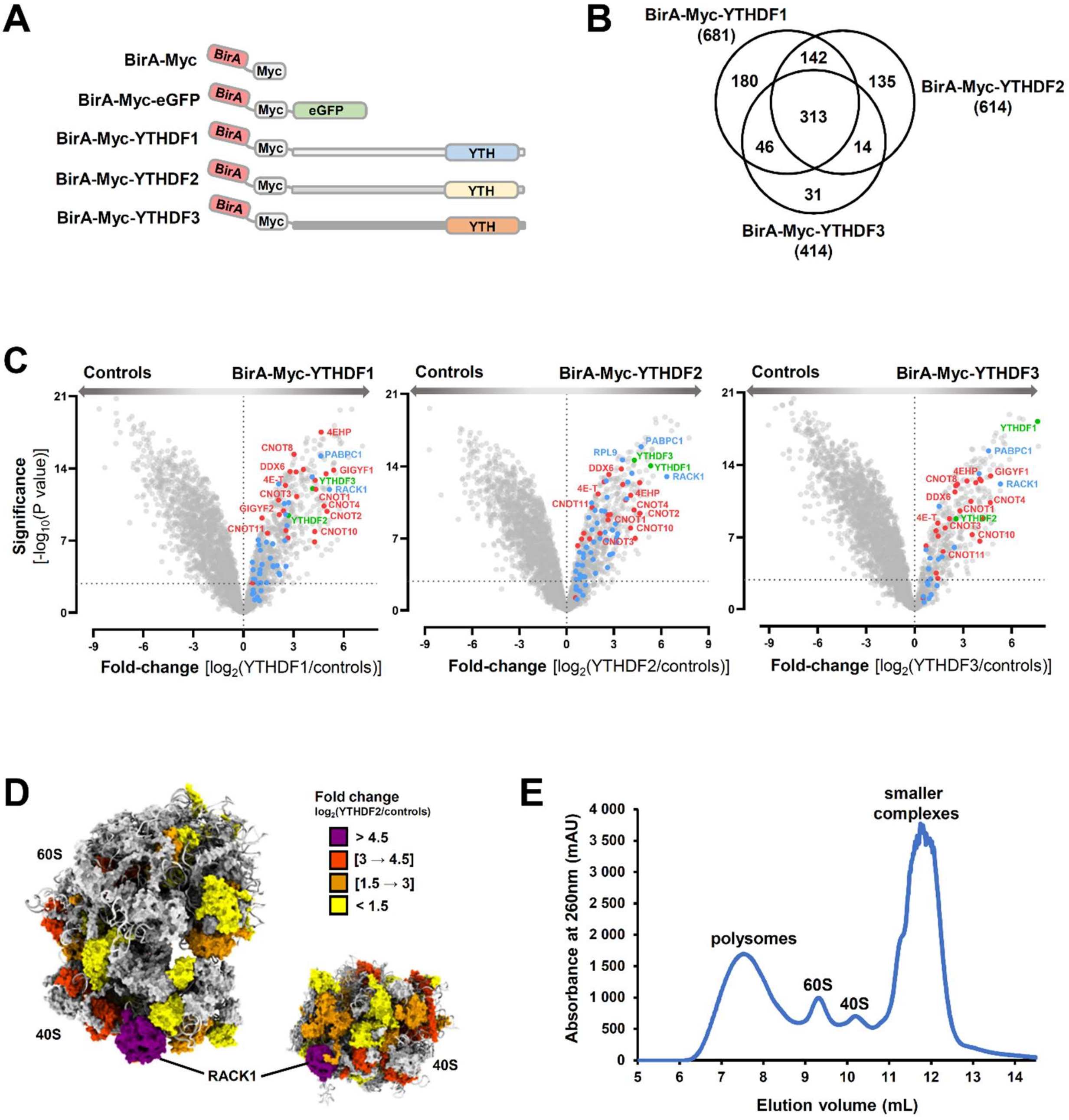


**Supplementary Figure S2 – BioID mapping of the YTHDF1-3 paralogs shows their proximity with ribosomes.** (**A**) Schematic representation of the BirA-Myc fused constructs used to generate the BioID datasets presented in panels B-C hereafter. BirA-Myc was individually fused to the N-terminus of YTHDF1, 2 and 3. (**B**) Venn diagram comparing the BioID datasets obtained with BirA-Myc-YTHDF1, 2 or 3. For each fusion, proximal proteins are listed in Supplementary tables 4-6. (**C**) Volcano plot showing proteins enriched in the BioID of BirA-Myc-YTHDF1, 2 and 3 over the control BioID samples (BirA-Myc alone + BirA-Myc-eGFP). The logarithmic fold-changes were plotted against negative logarithmic P values of a two-sided two samples t-test. A selection of silencing factors (red) and translation factors, including ribosomal proteins (blue) is indicated. The YTHDF paralogs are shown in green. (**D**) Ribosomal proteins were colored in the human ribosome (PDB: 6QZP) according to their logarithmic fold-changes calculated in the BirA-Myc-YTHDF2 BioID over the control BioID samples (BirA-Myc alone + BirA-Myc-fused eGFP). Ribosomal proteins in grey were not detected as high confident hits in the YTHDF2 BioID. (**E**) Chromatogram of the Ribo Mega-SEC fractionation shown in Figure 1F. The elution volume is indicated on the *x*-axis and the UV absorbance at 260 nm on the *y*-axis.

### Supplementary Figure S3

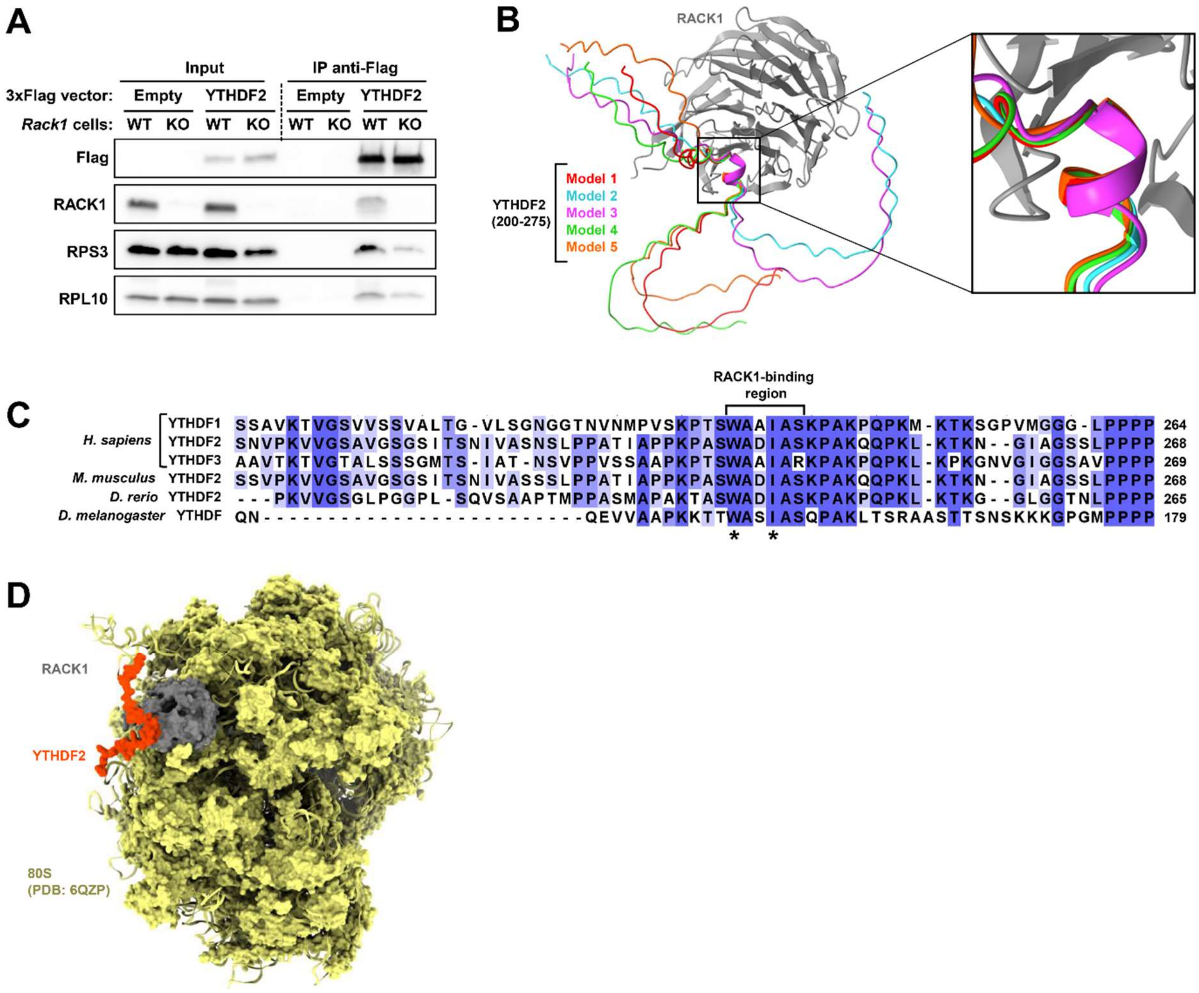


**Supplementary Figure S3 – RACK1 partially mediates the interaction between YTHDF2 and ribosomes**. (**A**) The interaction between YTHDF2 and ribosomal proteins is reduced in RACK1$^{KO}$ cells. Vectors expressing of 3xFlag-tagged YTHDF2, or 3xFlag as control (empty), were transfected into WT and RACK1$^{KO}$ HEK293T cells. Extracts were immunoprecipitated with anti-Flag antibody. Total lysates (input) and IP extracts were analyzed by Western blot (WB) with the indicated antibodies. (**B**) Superimposition of the five AlphaFold2-predicted models of the 200-275 segment of YTHDF2 bound to RACK1 (gray). The insert shows an enlarged view of the putative interface involving tyrosine 246 of RACK1 (Y246) stacked between tryptophan 239 (W239) and isoleucine 242 (I242). (**C**) Alignment of human YTHDF1, 2 and 3 with their homologous versions in *M. musculus, D. rerio* and *D. melanogaster*. The sequences were aligned with Clustal Omega using default parameters, and colored in Jalview by identity. Asterisks indicate the conservation of the W239 and 242 residues of human YTHDF2. (**D**) Superimposition of the AlphaFold2-predicted structure of the 200-275 segment of YTHDF2 (orange) bound to RACK1 (gray) on the structure of a human ribosome (yellow; PDB: 6QZP).

### Supplementary Figure S4

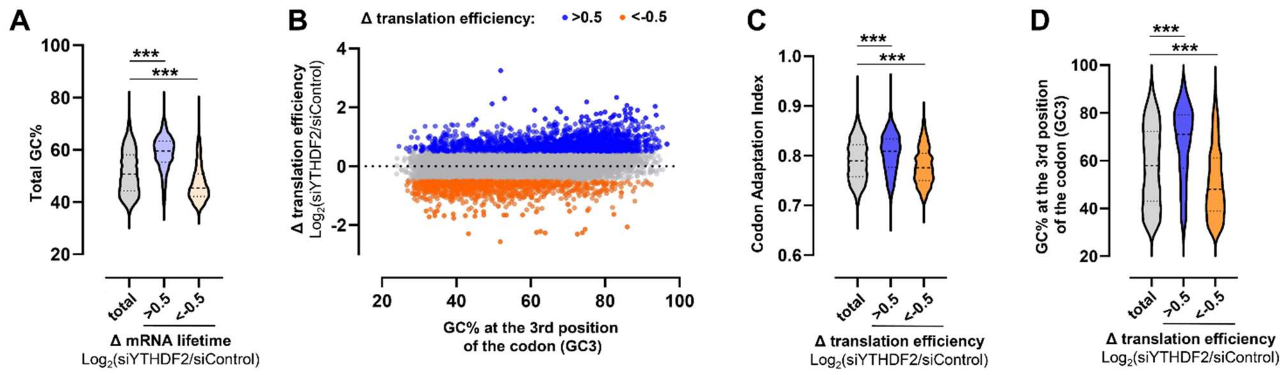


**Supplementary Figure S4 – YTHDF2 sensitivity among the transcriptome correlates with the GC content.** (**A**) Violin Plot showing the percentage of GC in the coding sequence (Total GC%) for each population of mRNAs shown in Figure 3A. P value was determined by a Kolmogorov-Smirnov test: (***) P < 0.0001. (**B**) Scatter plot representing changes in translation efficiency following YTHDF2 knockdown in Hela cells (publicly available data[10]) versus the percentage of GC at the third position of the codons of each transcript. A total of 8802 mRNAs are represented and colored according to the Δ translation efficiency (Log2(siYTHDF2/siControl)), including 1683 mRNAs with a Δ greater than 0.5, and 915 with a Δ lower than -0.5. (**C-D**) Violin Plot showing the Codon Adaptation Index (C) or the percentage of GC at the third position of the codons (D) for each population of mRNAs shown in (B). P value was determined by a Kolmogorov-Smirnov test: (***) P < 0.0001.

### Supplementary Figure S5

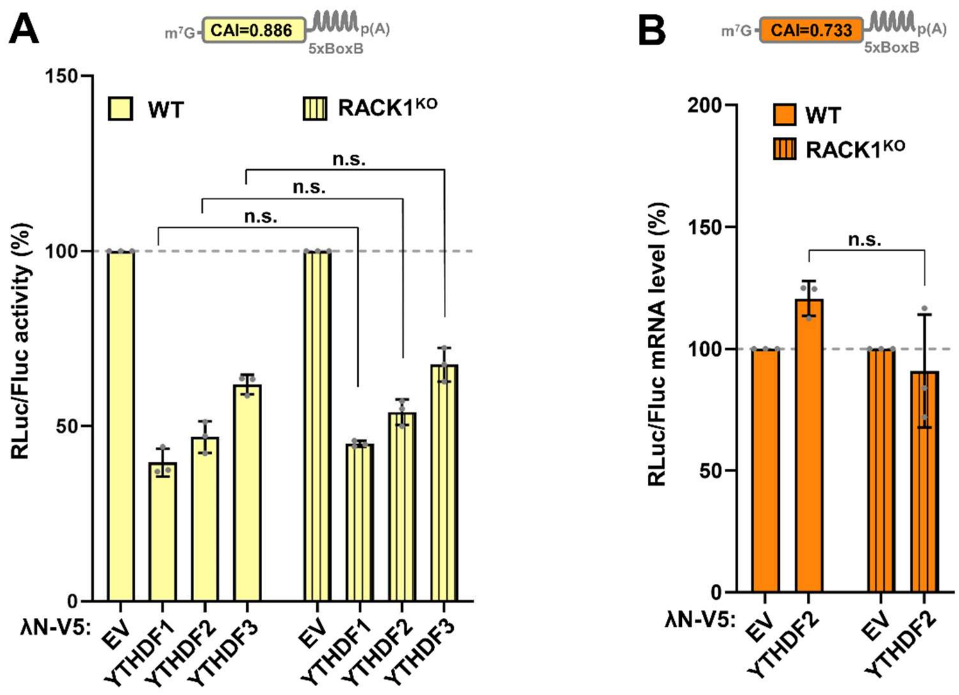


**Supplementary Figure S5 – Tethering the YTHDF proteins in RACK1$^{KO}$ cells.** (**A**) λN/BoxB tethering assay of the YTHDF protein on the RLuc$^{High}$ in RACK1$^{KO}$ cells. As in Figure 4E, WT and RACK1$^{KO}$ HEK293T cells were co-transfected with plasmids encoding λN-V5-YTHDF1-3, RLuc$^{High}$ and Firefly luciferase (FLuc). RLuc/FLuc activity values represent the mean ratio of RLuc luminescence normalized against FLuc expressed as percentage of the ratio of the cells transfected with the λN-V5 empty vector (EV). The mean values (±SD) from three independent experiments are shown and the P value was determined by two-tailed Student's t-test. (n.s., non-significant). (**B**) Measurements by RT-qPCR to estimate the RLuc$^{Low}$ and FLuc mRNA levels from the experiment described in Figure 4E.

### Supplementary Table S7. Oligonucleotides used in this study

| Fragments | Vectors | Sequences* (5'→3') | |
|---|---|---|---|
| XhoI-YTHDF1-NotI | pCI-Neo-3xFlag | Forward | AACTCGAGATGTCGGCCACCAGCGTGG |
| | | Reverse | AAGCGGCCGCTCATTGTTTGTTTCGACTCTG |
| XhoI-YTHDF2-NotI | pCI-Neo-3xFlag | Forward | AACTCGAGATGTCGGCCAGCAGCCTCTTG |
| | | Reverse | AAGCGGCCGCTTATTCTGAAGGAGTAGATCCAGA |
| XhoI-YTHDF2$^{\Delta 1\text{-}199}$-NotI | pCI-Neo-3xFlag/pCI-Neo-eGFP | Forward | AACTCGAGGCAAGCAATGTTCCAAAAGT |
| | | Reverse | AAGCGGCCGCTTATTCTGAAGGAGTAGATCCAGA |
| XhoI-YTHDF2$^{\Delta 1\text{-}275}$-NotI | pCI-Neo-3xFlag | Forward | AACTCGAGGGAACTTGGGATAACAAGGG |
| | | Reverse | AAGCGGCCGCTTATTCTGAAGGAGTAGATCCAGA |
| XhoI-YTHDF2$^{\Delta NT}$-NotI | pCI-Neo-3xFlag/pCI-Neo-eGFP | Forward | AACTCGAGATGCCCCACCCAGTGTTGGAGAAG |
| | | Reverse | AAGCGGCCGCTTATTCTGAAGGAGTAGATCCAGA |
| XhoI-YTHDF3-NotI | pCI-Neo-3xFlag | Forward | AACTCGAGATGTCAGCCACTAGCGTGG |
| | | Reverse | AAGCGGCCGCTTATTGTTTGTTTCTATTTC |
| EcoRI-YTHDF1-BamHI | pcDNA3.1 mycBioID | Forward | AAGAATTCATGTCGGCCACCAGCGTGG |
| | | Reverse | AAGGATCCTCATTGTTTGTTTCGACTCTG |
| EcoRI-YTHDF2-BamHI | pcDNA3.1 mycBioID | Forward | AAGAATTCATGTCGGCCAGCAGCCTCTTG |
| | | Reverse | AAGGATCCTTATTTCCCACGACCTTGACG |
| EcoRI-YTHDF2$^{\Delta NT}$-BamHI | pcDNA3.1 mycBioID | Forward | AAGAATTCATGCCCCACCCAGTGTTGGAGAAG |
| | | Reverse | AAGGATCCTTATTTCCCACGACCTTGACG |
| EcoRI-YTHDF3-BamHI | pcDNA3.1 mycBioID | Forward | AAGAATTCATGTCAGCCACTAGCGTGG |
| | | Reverse | AAGGATCCTTATTGTTTGTTTCTATTTC |
| EcoRI-eGFP-BamHI | pcDNA3.1 mycBioID | Forward | AAGAATTCATGGTGAGCAAGGGCGAGGA |
| | | Reverse | AAGGATCCTCACTTGTACAGCTCGTCCATGC |
| NheI-eGFP-XhoI | pCI-Neo | Forward | AAGCTAGCATGGTGAGCAAGGGCGAGGA |
| | | Reverse | AACTCGAGCTTGTACAGCTCGTCCATGC |
| SalI-RACK1-NotI | pGEX6p1 | Forward | AAAGTCGACTCACTGAGCAGATGACCCTTCG |
| | | Reverse | AAAGCGGCCGCCTAGCGTGTGCCAATGGTCAC |
| YTHDF2$^{\Delta 239\text{-}245}$ | pET-28c(+)-YTHDF2 | Forward | GCTCCTCCAAAACCAGCATCTCCTGCAAAACAGC |
| | | Reverse | GCTGTTTTGCAGGAGATGCTGGTTTTGGAGGAGC |
| YTHDF2$^{W239A/I242A}$ | pET-28c(+)-YTHDF2 | Forward | CATTGCTCCTCCAAAACCAGCATCTGCGGCTGATGCTGCTAGCAAGCCT |
| | | Reverse | AGGCTTGCTAGCAGCATCAGCCGCAGATGCTGGTTTTGGAGGAGCAATG |
| RACK1$^{Y246A}$ | pGEX6p1-RACK1 | Forward | TTCAGCCCTAACCGCGCCTGGCTGTGTGCTGC |
| | | Reverse | GCAGCACACAGCCAGGCGCGGTTAGGGCTGAA |
| RACK1$^{R36D/K38E}$ | pCI-Neo-3xFlag-RACK1 | Forward | CCCGGACATGATCCTCTCCGCCTCTGATGATGAGACCATCATCATG |
| | | Reverse | CATGATGATGGTCTCATCATCAGAGGCGGAGAGGATCATGTCCGGG |

*Restriction sites in the oligonucleotides are underlined.
